# Development of a Murine Model of Pyogenic Flexor Tenosynovitis

**DOI:** 10.1101/2020.02.07.925339

**Authors:** Bowen Qiu, Justin Cobb, Alayna Loiselle, Constantinos Ketonis

**Affiliations:** Department of Orthopaedics and Rehabilitation, Center for Musculoskeletal Research, University of Rochester Medical Center, Rochester, NY 14642

**Author notes:** Denotes equal contributions (Bowen Qiu, Justin Cobb). **Corresponding author:** Constantinos Ketonis, MD, PhD, Department of Orthopaedics, University of Rochester Medical Center, 601 Elmwood Drive Rochester, New York 14642.

**Keywords:** Murine model, pyogenic flexor tenosynovitis, Xen29

## Abstract

**Background:** To demonstrate the plausibility of a murine model of pyogenic flexor tenosynovitis.

**Methods:** 2μL of sterile PBS or bioluminescent Xen29 *Staphylococcus aureus* was administered to the tendon sheath of 36 male C57BL/6J mice. The infectious course was monitored by bioluminescence (BLI) signal via IVIS imaging and recording of weight change. The infected hind paws were harvested at four time points: 24 hours, 72 hours, 1 week and 2 weeks for histopathology using Alcian Blue hematoxylin staining. Two-way ANOVA with Sidak’s multiple comparison test was used for statistical analysis.

**Results:** The infected cohort displayed significantly elevated bioluminescent values, reductions in weight, and exhibited swelling of the infected digit throughout the course of infection. By day 7 most infected mice saw a substantial decrease in BLI signal intensity, however two infected mice exhibited persistent BLI intensity through day 14. Histopathology of the infected cohort showed tissue disorganization and the presence of a cellular infiltrate in and around the flexor tendon sheath.

**Conclusions:** A murine model of pyogenic flexor tenosynovitis is possible. Further optimization of the model offers an experimental platform for investigation of the pathophysiology of pyogenic flexor tenosynovitis.

**Clinical Relevance:** This animal model can be utilized in order to elucidate the basic molecular/cellular mechanisms of pyogenic flexor tenosynovitis while simultaneously evaluating novel therapeutic strategies.

## INTRODUCTION

Pyogenic flexor tenosynovitis (PFT), also known as septic or suppurative flexor tenosynovitis, is a severe closed-space bacterial infection that develops within the flexor tendon sheath of the hand. Kanavel first classified this condition more than a century ago as one of the most devastating infections in the upper extremities and described the four classic cardinal signs: pain with passive extension of the digit, symmetric digital swelling (fusiform swelling/sausage digit), tenderness along the flexor sheath, and a semi-flexed resting posture of the involved digit [1]. These infections usually follow a direct puncture and inoculation of the tendon sheath [2], although atraumatic hematogenous bacterial dispersion has also been reported [2]. The majority of PFTs are caused by *Staphylococcus aureus* [3], with isolation of methicillin-resistant *Staphylococcus aureus* (MRSA) in 29% of cases [3].

The presence of tendon sheath and bursa interconnections makes it possible for the infection to rapidly spread to other digits [2], and despite prompt treatment that typically includes surgical debridement and administration of antibiotics, PFT remains a great source of morbidity and disability [2]. Even if the infection is successfully eliminated, complications rates are unacceptably high [4]. Many patients will develop sequalae including stiffness, adhesions, tendon tears and tissue necrosis which in some cases can even lead to amputation of the infected digit [2]. Despite its high incidence and the degree of disability caused by PFT [5], the most efficacious treatment regimen still remains highly debated. Recent studies have demonstrated that the pathophysiology of this condition might be analogous to other closed space infections such as septic arthritis or periprosthetic infections[2]. Biofilm has been successfully formed on the tendon surface in a cadaveric model, which exemplifies a specific bacterial mechanism that could account for the challenges in treatment of PFT [4]. In light of these new observations, current treatment strategies are being reassessed.

Currently there is a scarcity of ideal animal models of PFT making this task difficult[6]. Such an animal model would significantly aid in the elucidation of the pathophysiology and bacterial mechanisms involved in PFT and would serve as an experimental platform for testing novel therapeutics for PFT. The murine hindpaw possesses deep and superficial flexor tendons that anatomically resemble zone 2 in the human hand and would constitute an excellent model for this infectious process. Moreover, mice are well studied, inexpensive, readily available and easy to house and handle and are thus a great translational model [7]. The aim of this study, therefore, is to establish a murine model of tenosynovitis which could aid in the development of an effective treatment strategy that is universally agreed upon. This could significantly benefit patients, while simultaneously decreasing the associated economic burden to the healthcare system.

## MATERIALS AND METHODS

### Animal Ethics

All procedures performed on animals were completed at the University of Rochester and were approved by the University Committee on Animal Research (UCAR: 2018-008). A strict adherence to the Guide for Care and Use of Laboratory Animals (IACUC) was implemented for this study.

### Experimental Outline

Thirty-six 14-week old C57BL/6J male mice received an intrasheath injection of either 1×10^7^ CFU *Xen29* or of 2μL PBS. Bioluminescent values of injected hindpaw were obtained on the following days: −1, 1, 2, 3, 4, 7, 10, and 14. Mice were weighed on following days −1, 1, 2, 3, 7, 10, and 14 and n=4 mice per group were harvested on days 1, 3, 7, and 14 for histological examination of the right hindpaw **(Figure 1)**.

**Figure 1:**
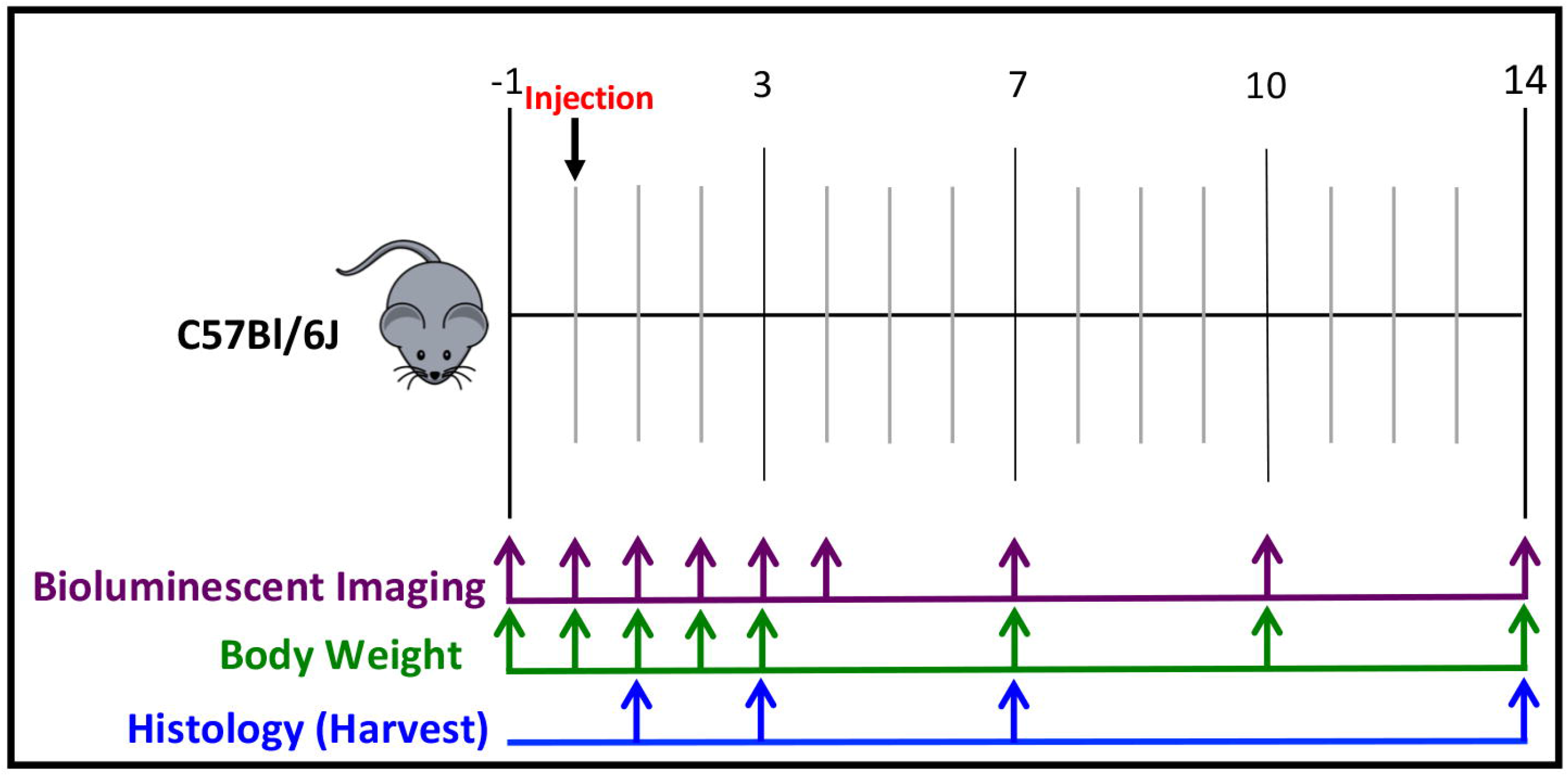
Experimental outline. 36 C57BL/6J mice had their baseline bioluminescence and weights taken on Day −1. On Day 0, the mice were either given a 2μL dose of 1×10^7^ CFU of Xen29 or a 2μL dose of 1X PBS by route of intrasheath injection. Mouse weight, bioluminescence, and clinical presentation were recorded throughout the infection time course. Mice were harvested on Day 1, Day 3, Day 7, and Day 14 for histological examination of the right hindpaw.

### Growth Conditions of S. aureus Xen29

Bioluminescent *Staphylococcus aureus* (Xen29; ATCC #12600) was streaked onto a Tryptic Soy Broth plate (TSB, Sigma-Aldrich #22092-500G) and incubated overnight at 37°C. For quality control purposes, the TSB plate was imaged to confirm positive detection of Xen29 bioluminescence. A single colony was obtained, placed in 3.0 mL of liquid Luria Bertani broth, and cultured for 23h at 37°C at 225 rpm. The next day, the cultures were centrifugated at 5,000 RPM for 5 minutes at 4°C. The resulting pellet was resuspended in 2.5 mL of sterile PBS (sPBS), with 100μL of culture set aside for OD_600nm_ measurements (OD_600nm_ = 0.589) and a ten-fold serial dilution for colony forming unit (CFU) calculations (1×10^7^ CFU per 2μL dose). Prior to the intrasheath injection, the bacterial suspension was mixed thoroughly.

### Animal Procedures: Intrasheath inoculation of S. aureus

To determine the feasibility and optimal parameters of intrasheath injection, mice were injected with varying volumes of Toluidine Blue (TB) dye (1μL, 2 μL, and 5μL). Successful injections were evaluated by the efficacy of TB dye localization to the tendon sheath with minimal detection of the TB dye in the surrounding tissue (**Supplemental Figure 1**). Thirty-six 14-week old C57BL/6J male mice were purchased from Jackson Laboratories (Stock #0664). Animals were housed on a 12-hour light/dark cycle and were ad libitum fed standard chow. Animals were administered pre-operative analgesia (1 mg/kg of Sustained-Release Buprenorphine), and were sedated with 100mg/kg Ketamine, via intraperitoneal injection. Postoperative pain monitoring was conducted for 3 days after each injection. A microsyringe (#7635-01, Hamilton Company, 7 Reno, Nevada) with a 30-gauge needle (#7803-07, Hamilton Company) was filled with 2μL of either 1×10^7^ CFU *Xen29* or sPBS. The needle was inserted into the medial aspect of the third digit on the right hindpaw between the proximal interphalangeal joint (PIP) and the metatarsophalangeal (MTP) joint, utilizing forceps for stabilization of the third digit. This digit was chosen because it is the longest and largest digit so as to provide the highest likelihood of entering the sheath.

### In vivo detection of Xen29 infection propagation

Prior to bioluminescent detection, animals were placed in a chamber and given 2.5% isoflurane (Fluriso, VetOne) to induce temporary anesthesia. A single mouse was taken from the isoflurane chamber and placed in a prone position, with the right hindpaw stabilized, in the IVIS® Spectrum (PerkinElmer) imaging chamber. 1.5% isoflurane was applied via a vaporizer during the imaging process. A region of interest was established on a sPBS injected mouse, beginning at the medial aspect of the hindpaw and encompassing all of the digits. This region of interest was subsequently applied to the remainder of the animals and bioluminescence was quantified as Radiance (photons/s/cm^2^/sr). Bioluminescent Imaging was conducted on days: −1, 1, 2, 3, 4, 7, 10, and 14.

### Histological Characterization

Animals were euthanized at 24h, 72h, 1 week, and 2 weeks post-infection via CO_2_ overdose and cervical dislocation. The hind paws were harvested proximally to the palm and placed into 10% neutral buffered formalin (NBF) for 72h at room temperature with rocking. The hind paws were then washed with PBS and dH_2_O and placed in 14% EDTA decalcification (Webb-Jee) solution for 1 week, with daily changes of the solution. Following routine paraffin processing, 5μm axial sections were obtained and the sections were subsequently stained with Alcian Blue Hematoxylin (ABHOG) to visualize changes in tissue morphology.

### Statistical Analysis

Statistical significance for the bioluminescent values and percent weight change values between the infected cohort and the control group at each time point were calculated using two-way ANOVA testing with post-hoc Sidak’s multiple comparison testing.

## RESULTS

### Temporal bioluminescence monitoring

*In vivo* Xen29 bioluminescent signal detection allows for longitudinal monitoring of persistent *S. aureus* infection.

IVIS imaging on post-injection day 1-14 demonstrate good localization of infection to the middle digit without spread to the palm or other digits indicating establishment of isolated infection within the tendon sheath **(Figure 2).** Prior to injection, baseline bioluminescence values between the two cohorts were not statistically different. The infected group of animals displayed a significant increase in bioluminescence (p < 0.0001) for the first 2 days after infection **(Figure 2B)**, relative to sPBS. On day 3 post-injection, the majority of S. aureus injected animals retained a strong bioluminescence signal, although the mean BLI signal was not significantly different than sPBS injected digits **(Figure 2B).** After post-injection day 3, most *S. aureus* injected animals exhibited a substantial decrease in BLI intensity, with signal intensity approaching that of sPBS injected controls (this group was stratified as ‘Subacute infection’ for histological analysis). However, two mice retained a robust bioluminescence signal until the experimental endpoint **(Figure 2C)**, (these mice were stratified as ‘Active infection’ for histological analysis.

**Figure 2:**
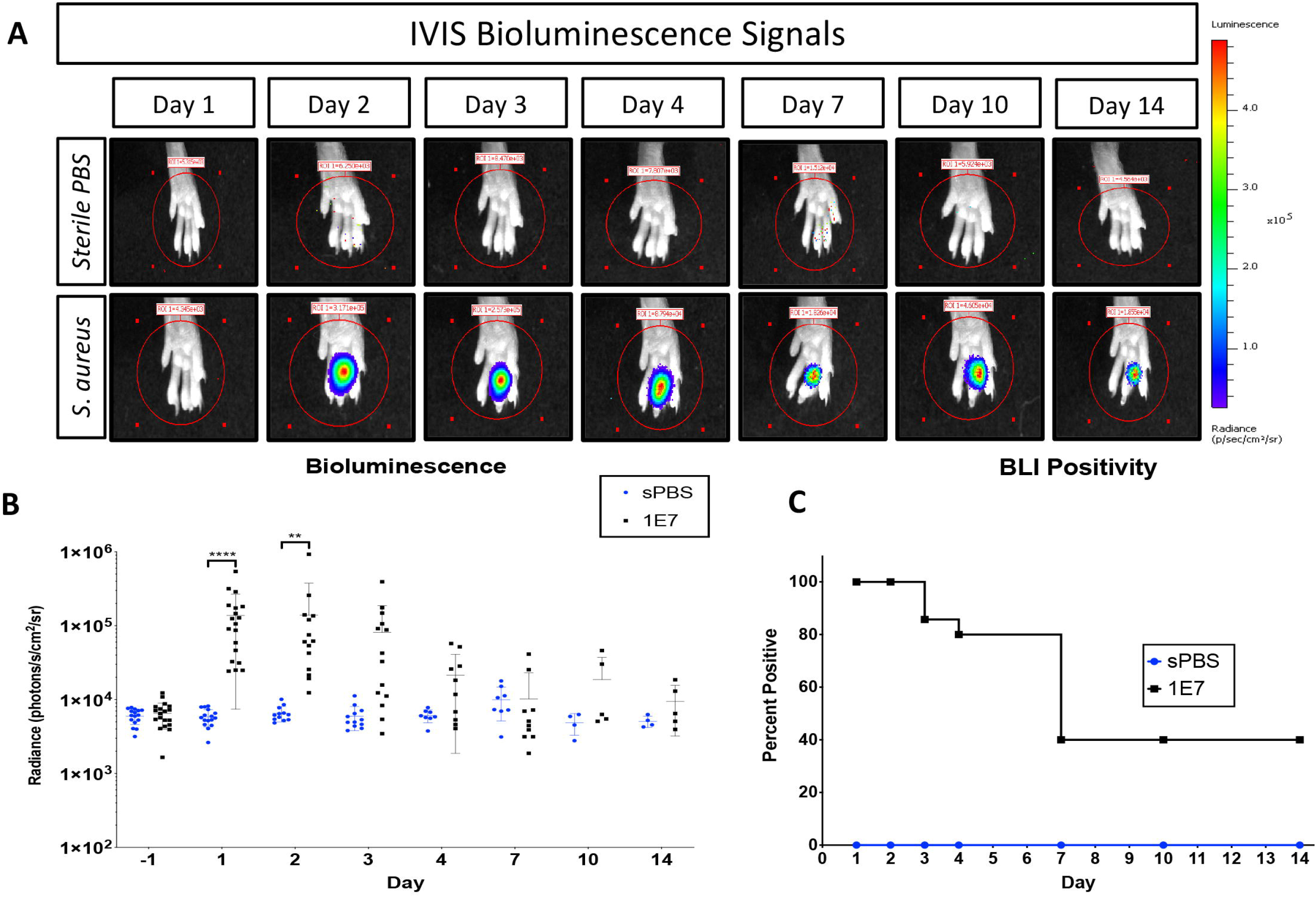
A.) In vivo bioluminescence obtained using the IVIS imaging system. Signal is present and localized to the middle digit as compared to controls. B.) In vivo bioluminescent detection of Xen29 in the murine tendon sheath. Units are expressed as Radiance (photons/s/cm^2^/sr). Ordinary two-way ANOVA with Sidak’s multiple comparison test. **** = p < 0.0001, ** = p < 0.001. C.) Survival curve illustrating the percentage of positive BLI signal throughout the experimental time course.

### Clinical markers of infection

Daily weights were also measured for all animals as weight loss is an indication of persistent infection. The weights obtained on Day −1 were normalized to 100% and reported as percent body weight lost during the infection process. Infected animals had significantly lower weights as compared to the control animals on day 2 through 10 (p < 0.05) **(Figure 3)**. Gross observation of the infected digits for all animals showed development of fusiform swelling which persisted even with the loss of bioluminescence signal **(Figure 4)**. This feature was notably absent in animals in the control group that received an intra-sheath injection with saline.

**Figure 3:**
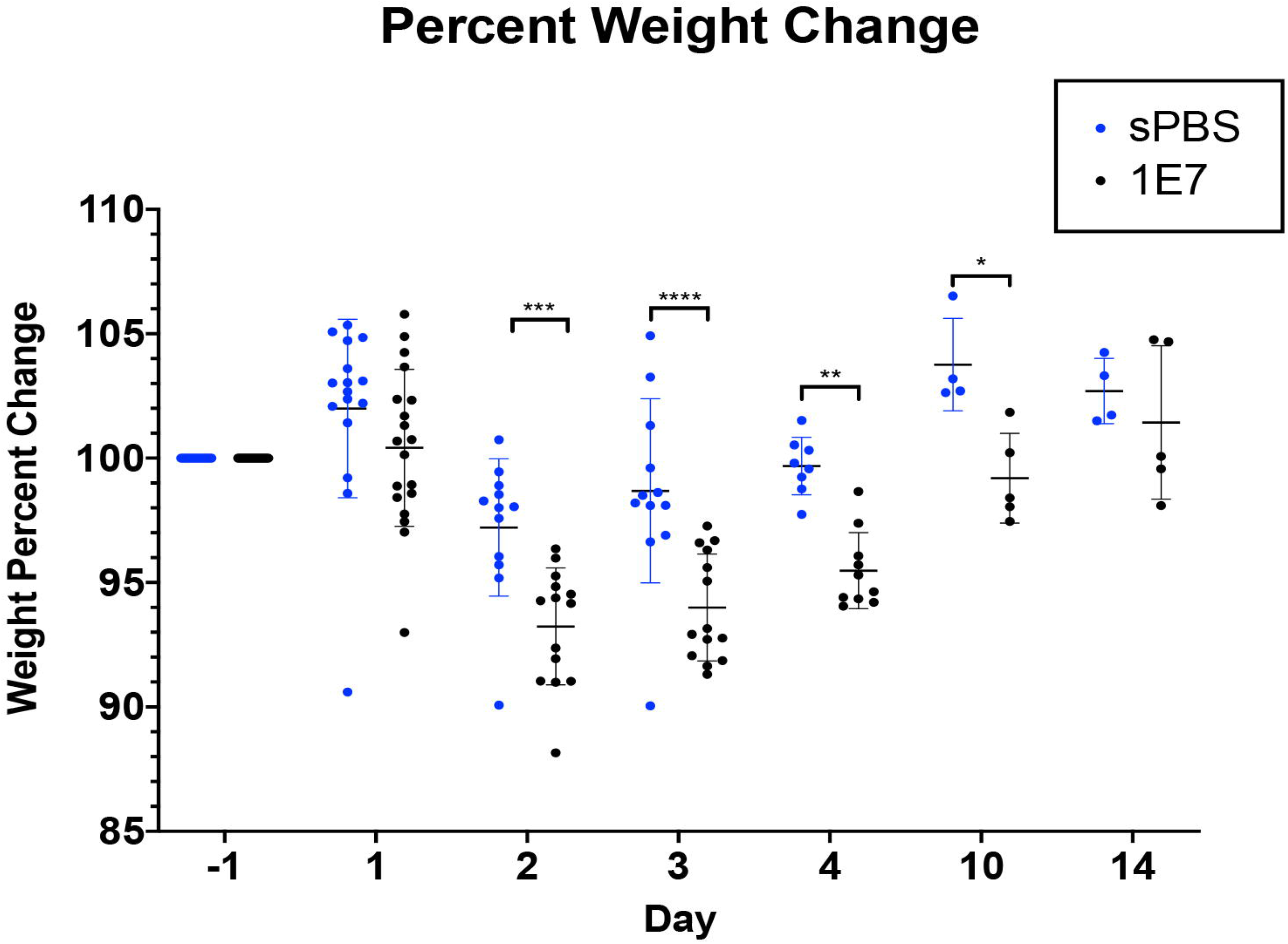
Weight loss expressed as percentage of weight lost post-intrasheath injection. Day −1 weight values were used to normalize percent weight increase or decrease for each individual mouse. Ordinary two-way ANOVA with Sidak’s multiple comparison test. ****= p < 0.0001, ***= p < 0.0005, **= p < 0.005, *= p < 0.05

**Figure 4:**
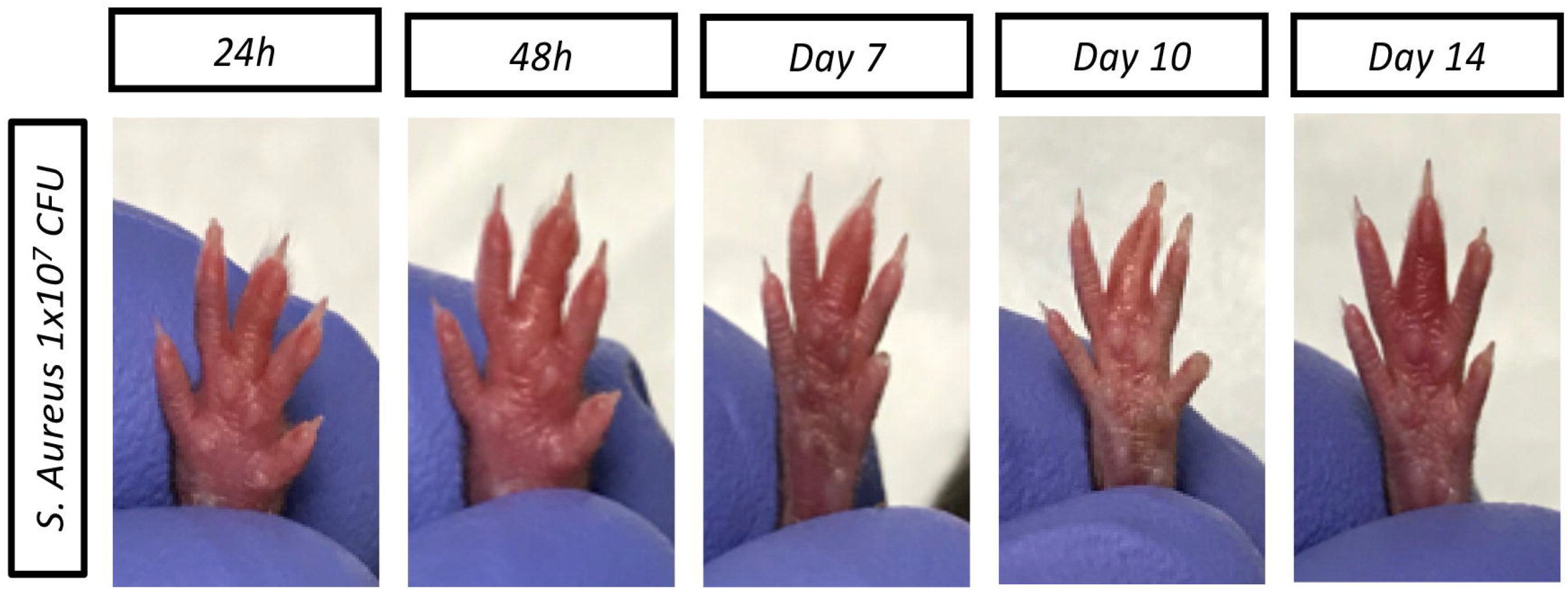
Gross morphological characterization of intrasheath injection of Xen29 at five timepoints: 24 hours, 48 hours, 72 hours, 1 week, and 2 weeks post-injection. At each time point there was a clear indication of swelling of the infected digit. This swelling could be the murine analog to fusiform swelling, which is a clinical diagnostic observation for PFT in humans. Photos were taken at the specified time points.

### Histological characterization exhibits cellular infiltration and tissue disorganization

For histological analysis the animals were divided in 3 different groups: control group, active infection (animals with persistently high bioluminescent signal) and subacute infection (animals with clinical signs of infection without significant bioluminescent signal). The control group **(Figure 5, top row)** demonstrated preserved synovial space between FDS and FDP tendons with no evidence of fibrosis throughout Day 14. In contrast, both active and subacute groups **(Figure 5, middle and bottom row)** exhibit dense cellular infiltration in and around the tendon sheath. Pronounced thickening of the sheath itself is observed with infiltration and collapse of the synovial space. Tendon morphology showed a gradual distortion and deterioration as infection progressed from Day 3 to Day 14. **(Figure 5)**. Also notable is the significant thickening of the floor of the tendon sheath when compared to the control group (**arrows**). The gray, fibrous features have the typical appearance of hypertrophic scars and are more pronounced at the latter time points of 1 and 2 weeks **(Figure 5, column 3 and 4)**. A subacute or resolving infection resulted in a more structurally disorganized tendon phenotype, than was evident in the sustained infection. Two animals in the infected cohort displayed a phenotype that resembled cortical bone formation, potentially suggesting a transition in the infection to osteomyelitis **(Supplemental Figure 2)**.

**Figure 5:**
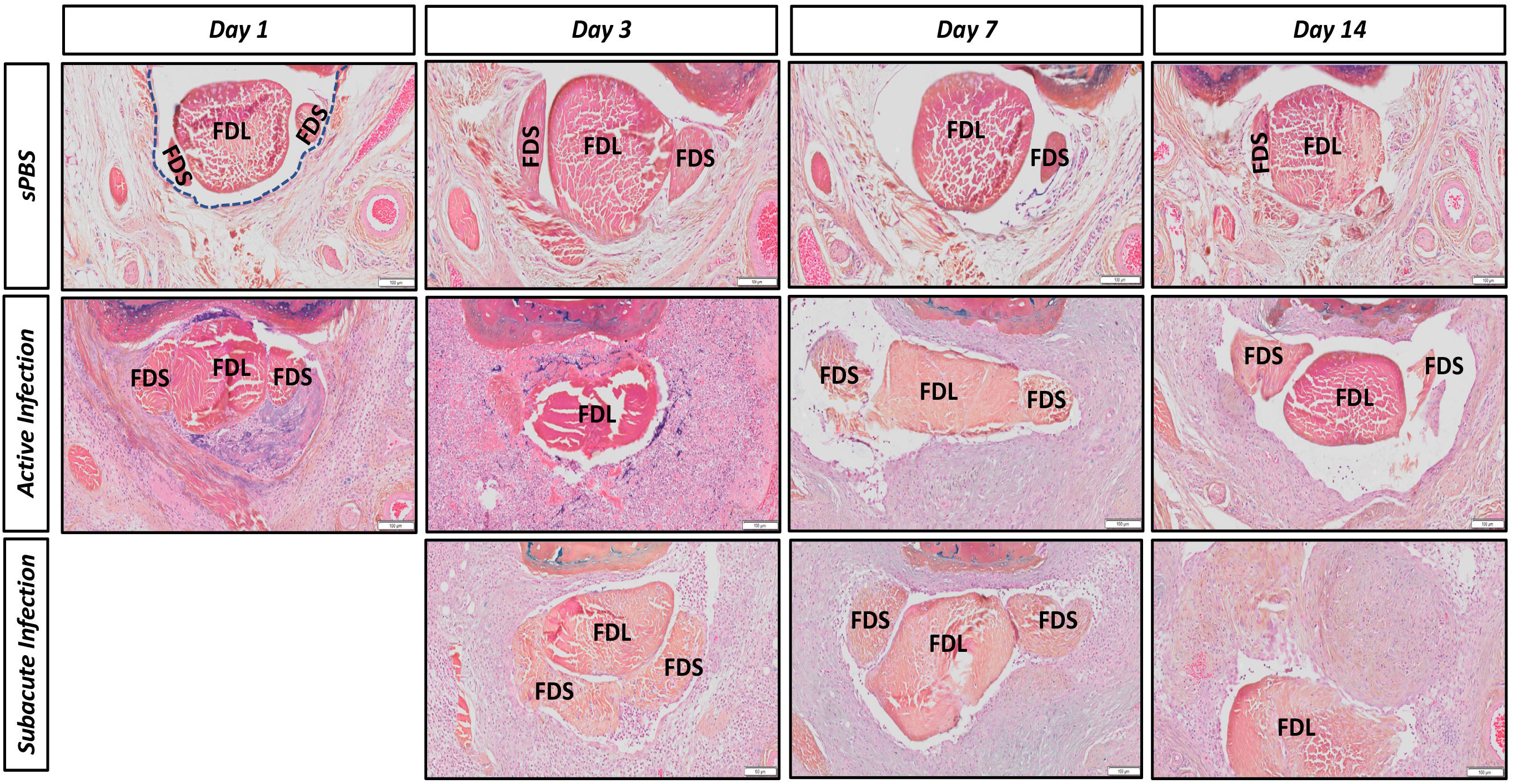
Alcian Blue Hematoxylin/Orange G stain. Sustained infection indicates the presence of bioluminescent signal at the time of harvest. Cleared infection indicates the absence of bioluminescent signal at the time of harvest. Arrows indicate excess tissue deposition. Images were taken at 8x magnification.

## DISCUSSION

Pyogenic flexor tenosynovitis is a devastating hand infection. However, despite its heavy disease burden, there remains no animal model that accurately recapitulates this disease process. In this study we demonstrated the feasibility of developing a murine model of pyogenic flexor tenosynovitis following direct inoculation of the flexor sheath. The infected animals followed a clinical course typical for flexor tendon infections, exhibiting fusiform swelling, redness, and scarring and displayed appropriate bioluminescence signal through the course of the disease, while histopathologic analysis of the infected digit demonstrated changes consistent with flexor tenosynovitis. [1]

The concentration and volume of *S. aureus* inoculate has greatly varied in the orthopedic implant and biofilm literature, with dose dependent effects on the host immune/inflammatory response [8]. In our experiments we found that concentrations too low led to premature clearance of infection or no infection at all, while concentrations too high led to frank necrosis of the digit (data not shown). Optimal inoculate concentrations are disease specific, and even though other studies using Xen29 demonstrate successful establishment of infection with concentrations on the magnitude of 10^4^ CFU [9], intra-sheath inoculation in this model required 1×10^7^ CFU for reproducible colonization. It should be noted however that mice are not a natural host for human clinical *S. aureus* and the concentrations needed to induce infection can be significantly higher [10, 11].

Bacterial bioluminescence is a common method to track propagation of bacteria with lux operons, such as Xen29[12]. Previous studies utilizing Xen29 note bioluminescence signal peaking at around post-infection day 3, and remains elevated until post-infection day 7, before beginning to dissipate; a timeline that matches well with the pattern observed in our study.[9, 13] Using bioluminescence as a proxy for the presence of bacteria, we found that infected mice had elevated signals throughout the first 7 days, at which point all but two of these mice lost signal. Clinically, these mice exhibited the same signs of flexor tendon infections and weight loss as those with persistent signals. We believe that these data demonstrate two possible outcomes of untreated flexor tendon infections: active infection and subacute infection. While the active infection group meets our expectations for typical infection models: persistent bioluminescence signal, clinical changes consistent with infectious process, and inflammatory hypercellularity, cessation of BLI signal can be attributed to multiple causes and does not necessarily mean that the infection was cleared. Previous studies have suggested that this may be due to a metabolic shift in the bacteria into a dormant phenotype like those found in biofilm, or that the bacterial chromosome has lost the *Photorhabdus luminescens* lux operon required for bioluminescence. Alternatively, this might suggest that there was a lack of substrate within the tendon sheath to produce the protein, or the bacteria have been eradicated [9, 14, 15]. In addition, low bioluminescence from an intact protein may also not be observable. Daghighi *et al*., showed in their study that despite loss of Xen29 signal in 7 animals, 4 out of 7 remained culture positive [16], suggesting that while a positive BLI signal indicates that that the lux operon is present – and by extension Xen29, the loss of BLI signal does not necessarily mean total clearance of infection. We do think that given the clinical outcomes of fusiform swelling and continued weight loss into day 10, that these mice were likely at a subacute level of infection.

Our clinical findings mirror that of infection, with weight as a main outcome measured. Infected mice weighed significantly less than the controls and was inversely associated with BLI signal intensity. Maximum weight deficit was observed at the same time as the maximum signal intensity, consistent with similar studies that track weight loss, infection, and bioluminescence signaling [13, 17, 18]. At 7 days, though 3 of the animal lost bioluminescence signals, weight remained significantly lower until day 10, suggesting that either infection was not completely eradicated at that time, or that the disease burden slowed recovery.

Histologically, we observed the presence of a cellular infiltrate in the tendon sheath and the surrounding tissue as a primary effect of Xen29 intrasheath injection. There was also persistence of tissue damage after cessation of the signal implying a dysregulated tissue healing processes as evident by the formation of potential hypertrophic scarring in the latter time points. This dysregulation may have been established by the expression of various Xen29 virulence factors throughout the course of acute infection. Staphylococcal infections have the potential to induce immunomodulatory changes to the infiltrating innate immune cells through expression of various virulence factors [9, 10]. Examination of the contribution of *S. aureus* virulence factors versus dysregulated tissue repair mechanisms could provide insights into the processes that contribute the most to the damage to the tissue. These virulence factors may result in an increased and/or sustained proinflammatory cellular environment, resulting in the degradation of tissue structure/tendon morphology. The histological findings of scar formation collapse of intra-sheath space and thickening of the sheath itself could account for the typical clinical picture and sequala of adhesions, stiffness and disfunction following PFT.

Elucidation of the cellular infiltrate populations could provide clues to the pathogenesis of murine PFT, especially by monitoring any changes to this population when antibiotics and/or corticosteroids are administered[19]. A description of the cytokine environment at varying time points can provide a time scale for length of proinflammatory environment. Coupling the cytokine expression with transcription factors (i.e. NF-KB) [20] or inflammatory proteins (i.e. C-reactive protein)[21] can offer a more detailed description of the molecular mechanisms of PFT that could potentially identify novel therapeutic targets in order to reduce scarring and stiffness. Finally, a murine model of PFT allows for testing the efficiency of experimental therapies, in addition to incorporating more sophisticated techniques for the experimental endpoints.

## Acknowledgements

We would like to thank the Biomechanics, Biomaterials and Multi-Modal Tissue Imaging (BBMTI) and the Histology, Biochemistry, and Molecular Imaging (HBMI) Cores in the Center for Musculoskeletal Research at the University of Rochester Medical Center for their technical assistance. The BBMTI and HBMI Cores are supported in part by NIH/ NIAMS P30 AR069655. This work was supported in by part a P30 Pilot Grant to CK.

## Supplemental Figures

**Supplemental Figure 1:** Toluidene blue injection demonstrating successful intra-sheath inoculation with little to no dye observed subcutaneous or within other structures of the mouse digit

**Supplemental Figure 2:** Histopathology of a limited number of animals (n=2) displayed cortical bone formation, suggesting the infection transitioned to osteomyelitis. Images were taken at 4x magnification.

