## Supplementary figures and images for "Development of a Murine Model of Pyogenic Flexor Tenosynovitis"

### Supplemental Figure 1

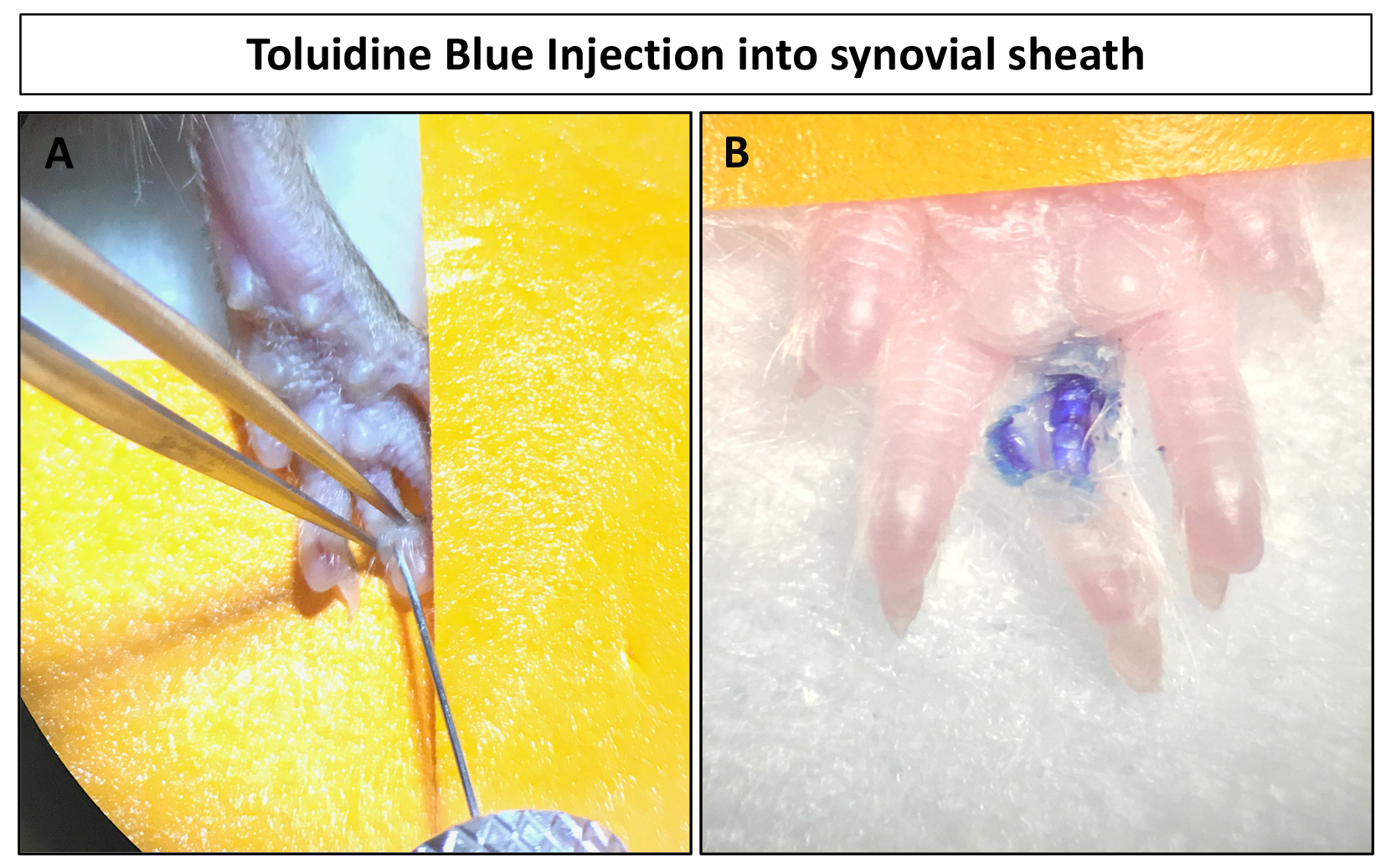

### Supplemental Figure 2

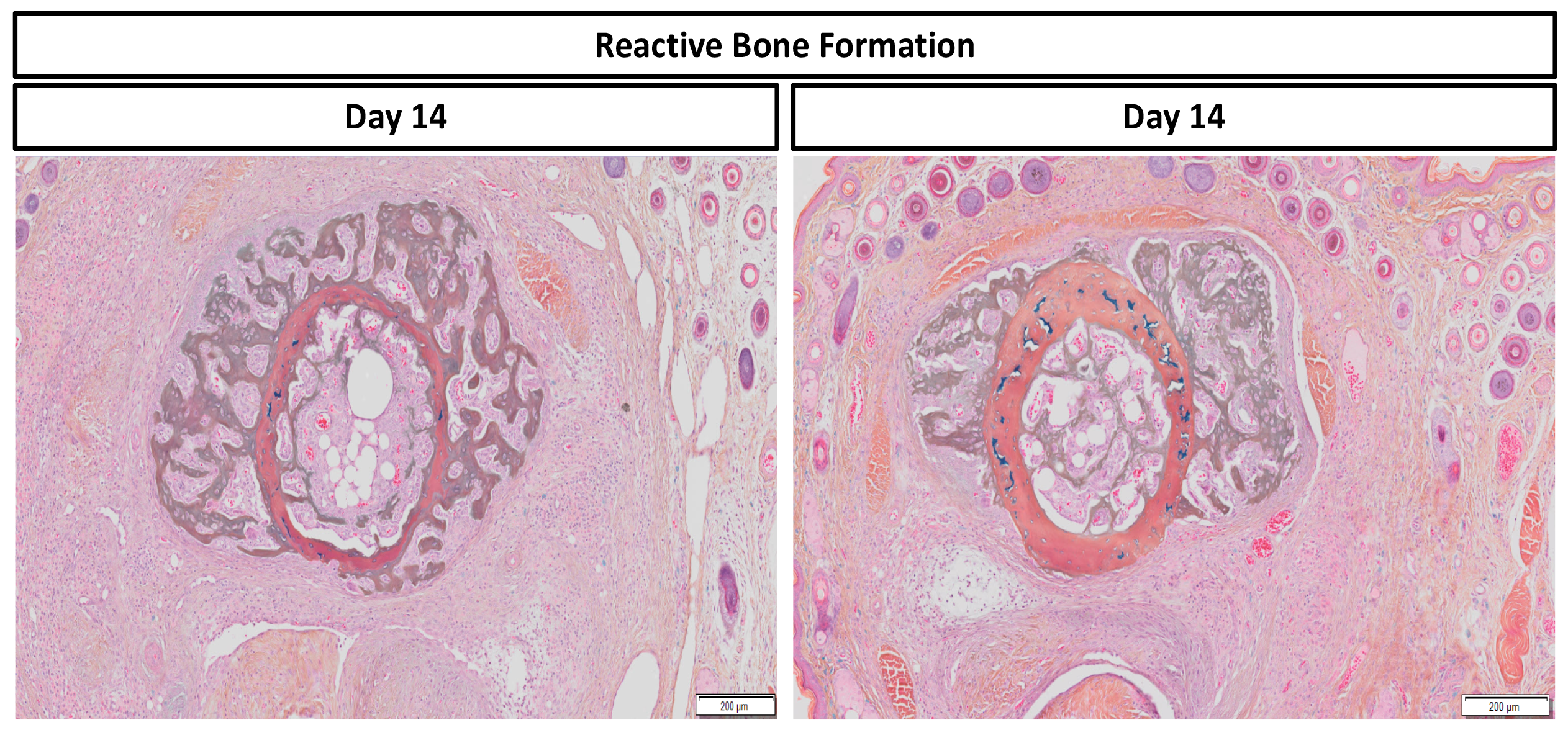
